# Metafounders are Fst fixation indices and reduce bias in single step genomic evaluations

**DOI:** 10.1101/083675

**Authors:** Carolina Andrea Garcia-Baccino, Andres Legarra, Ole F Christensen, Ignacy Misztal, Ivan Pocrnic, Zulma G Vitezica, Rodolfo J.C. Cantet

## Abstract

**BACKGROUND:** Metafounders are pseudo-individuals that condense the genetic heterozygosity and relationships within and across base pedigree populations, i.e. ancestral populations. This work addresses estimation and usefulness of metafounder relationships in Single Step GBLUP.

**RESULTS:** We show that the ancestral relationship parameters are proportional to standardized covariances of base allelic frequencies across populations, like Fst fixation indexes. These covariances of base allelic frequencies can be estimated from marker genotypes of related recent individuals, and pedigree. Simple methods for estimation include naïve computation of allele frequencies from marker genotypes or a method of moments equating average pedigree-based and marker-based relationships. Complex methods include generalized least squares or maximum likelihood based on pedigree relationships. To our knowledge, methods to infer F_st_ coefficients and F_st_ differentiation have not been developed for related populations.

A compatible genomic relationship matrix constructed as a crossproduct of {−1,0,1} codes, and equivalent (up to scale factors) to an identity by state relationship matrix at the markers, is derived. Using a simulation with a single population under selection, in which only males and youngest animals were genotyped, we observed that generalized least squares or maximum likelihood gave accurate and unbiased estimates of the ancestral relationship parameter (true value: 0.40) whereas the other two (naïve and method of moments) were biased (estimates of 0.43 and 0.35). We also observed that genomic evaluation by Single Step GBLUP using metafounders was less biased in terms of accurate genetic trend (0.01 instead of 0.12 bias), slightly overdispersed (0.94 instead of 0.99) and as accurate (0.74) than the regular Single Step GBLUP. Single Step GBLUP using metafounders also provided consistent estimates of heritability.

**CONCLUSIONS:** Estimation of metafounder relationship can be achieved using BLUP-like methods with pedigree and markers. Inclusion of metafounder relationships improves bias of genomic predictions with no loss in accuracy.

## BACGROUND

The concept of metafounders gives a coherent framework for a comprehensive theory of genomic evaluation [1]. Genomic evaluation in agricultural species often implies partially genotyped populations, i.e. some individuals are genotyped, others are not, and phenotypes may be recorded in either of the two subsets. An integrated solution called Single Step has been proposed [2–4]. This solution proposes an integrated relationship matrix

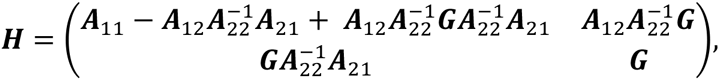

with inverse

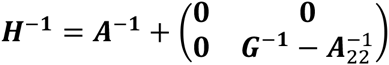

where ***G*** is the genomic relationship matrix, ***A*** is the pedigree-based relationship matrix, and matrices ***A***_11_, ***A***_2_, ***A***_21_, ***A***_22_ are submatrices of ***A*** with labels 1 and 2 denoting non-genotyped and genotyped individuals, respectively.

Because genotyped animals are not a random sample from the analyzed populations (they are younger or selected), it was quickly acknowledged that a proper analysis requires specifying different means for genotyped and non-genotyped individuals for the trait under consideration. These different means can be considered as parameters of the model, which are either fixed [4] or random [5,6]. In the latter case, the random variables induce covariances across individuals, a situation that is referred to as “compatibility” of genomic and pedigree relationships. In fact, compatibility implies comparability of the average breeding value of the base population and of the genetic variance [7] across the different measures of relationships.

Numerically, the problem shows up as follows. The formulae for matrix **H** and its inverse contain (**G** − **A**_22_) and 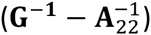 (assuming **G** is full rank), respectively. This suggests that if **G** and **A**_22_ are too different, biases may appear.

Genomic relationships are usually computed in one of two manners: the “crossproducts” [8] or the “corrected identity by state (IBS)” [9]. Both depend critically on assumed *base allelic frequencies* (Toro et al., 2011). However, for most purposes allelic frequencies are not of interest *per se* and can be treated as nuisance parameters to be marginalized. Christensen [10] achieved an algebraic integration of allele frequencies, leading to a very simple covariance structure with allele frequencies in genomic relationships fixed at 0.5 (e.g., using genotypes coded as {−1,0,1} in the crossproducts) and a parameter called *γ* which describes the relationships across founders i.e. 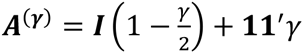 in the base population. A second parameter in Christensen’s marginalisation is *s*, which is a counterpart of the heterozygosity of the markers at the base population. Therefore, instead of inferring (thousands of) base allelic frequencies, inference can be based on two simple parameters *γ* and *s.* Both can be estimated maximizing the likelihood of observed genotypes. Also this considers the fact that pedigree depth is arbitrary and mostly based on historical availability of records.

Legarra *et al.* [1] showed the equivalence of Christensen’s ideas to metafounders: pseudo-individuals that simultaneously consider three ideas: (a) separate means for each base population [4,11], (b) randomness of these separate means [5] and (c) the propagation of the randomness of these means to the progeny [10], while accommodating several populations with complex crosses e.g. [12]. Legarra *et al.* [1] also generalized one relationship across founders (scalar *γ*) to several relationships across founders in the pedigree, i.e. ancestral relationships (matrix ***Γ***), and suggested simple methods to estimate them. However, the performance of their model, both for estimation of ancestral relationships and for genomic evaluation, has not been tested so far.

This work has two objectives. The first one is to delve into the structure of the metafounder approach to find an alternative parameterization and estimation of the ancestral relationships. By doing so we find that ancestral relationships are generalizations of Wright’s F_st_ fixation index. The second goal is to test, by simulation, (i) methods to estimate ancestral relationship parameters, (ii) the quality of genomic predictions using metafounders and (iii) the quality of variance component estimation. For the second goal, the simulated population is undergoing selection and with a complete pedigree partially genotyped.

## METHODS

### Relationship between metafounders and allelic frequencies at the base

#### Single population

Let ***M*** be a matrix of genotypes coded as gene content, i.e. {0,1,2} and the genomic relationship matrix ***G*** = (***M*** − ***J***)(***M*** − ***J***)’/*s* with ***J*** a matrix of 1’s, with reference alleles taken at random so that for a random locus the expected allelic frequency *p* is 0.5. [10]. In other words, the matrix ***Z*** = (***M*** − ***J***) contains values of {−1,0,1} for each genotype. In a single population, let *γ* be a relationship coefficient across pedigree founders or, equivalently the self-relationship of the metafounder [1,10]. Parameter *γ* is the relationship coefficient among the founders of a population, so that ***G*** = (***M*** − ***J***)(***M*** − ***J***)’/*s* is most likely given the observed pedigree. This relationship *γ* is relative to a population with maximum heterozygosity and it is analogous to an Fst fixation index. The parameter *s* is a measure of maximum heterozygosity in the population.

Christensen (2012) estimated the two parameters, *γ* and *s* using maximum likelihood, whereas Legarra et al. (2015) suggested methods of moments. Closer inspection of Appendix A in Christensen (2012) leads to the following developments (see supplementary material for more details).

The parameter *γ* is such that 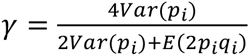 with *p*_*i*_ = 1 − *q*_*i*_ the allelic frequency at a random locus *i*. The parameter *s* = *n*(2*Var*(*p*_*i*_) + *E*(2*p*_*i*_*q*_*i*_)) with *n* being the number of markers. However, *E*(2*p*_*i*_*q*_*i*_) = 2*E*(*p*_*i*_)*E*(*q*_*i*_) − 2*Var*(*p*_*i*_) = 0.5 − 2*Var*(*p*_*i*_), where it was used that if alleles are labelled at random across loci then *E*(*p*_*i*_) = *E*(*q*_*i*_) = 0.5. From this it follows that 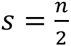 and the genomic relationship matrix is ***G*** = 2(***M*** − ***J***)(***M*** − ***J***)’/*n*. Interestingly, this matrix is similar to a matrix of IBS relationships, that can be written as ***G***_*IBS*_ *=* (***M*** − ***J***)(***M*** − ***J***)’/n + **11**′, so that 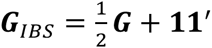. (See proof in the supplementary Material).

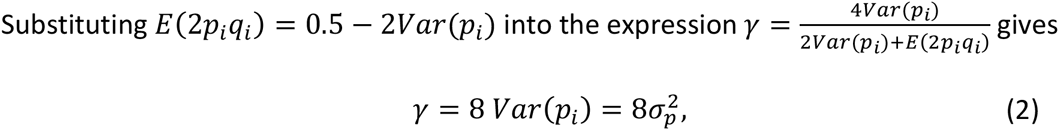

so that *γ* for a single population is eight times the variance of allelic frequencies at the base population (this variance was described by Cockerham [13]). These equalities were not described in Christensen [10]. We stress that 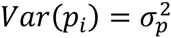 to imply that 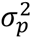 (and *γ*) is a parameter, the variance of allelic frequencies [10,14–16]. On the other hand, *s* can be seen as the heterozygosity in the case that all markers had an allelic frequency of 0.5.

#### Multiple populations

In an analogous manner, the relationship across two metafounders *b* and *b*′ is

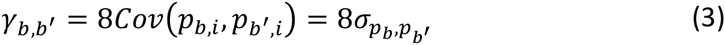

i.e., the covariance across loci between allelic frequencies of two populations *b* and *b*′. This is almost tautological: the relationship is the covariance across gene contents at a locus, here applied for populations. Christensen et al. (2015) show this in Appendix A, somehow implicitly. Cockerham [13] and Robertson [17] interpret 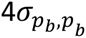 as the coancestry across two populations and Fariello et al. [18] use 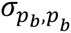, to describe the divergence of populations. There are several measures of genetic distance between populations (e.g. [19]), and most of them contain a term related, implicitly or explicitly, to 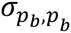,. In particular, the average square of the Euclidean distance can be written as 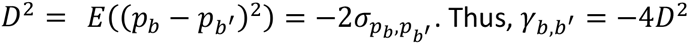.

### Estimation

#### Estimation in a single population

Estimation of *s* is trivial, it is simply half the number of markers. Parameter *γ* is proportional to the variance of allele frequencies. If base population individuals were genotyped, computing allele frequencies and estimating *γ* is trivial. In the next section we propose methods when this is not the case, i.e. genotyped individuals are related and perhaps several generations away from the base.

##### 1-Assuming no pedigree structure

<u>NAIVE</u>: The simplest model assumes that genotyped individuals are unrelated and constitute the base population. For locus *i*, let *m*_*i*_ be a vector of gene contents in the form {0,1,2}, defined as before. The mean of this vector is *μ*_*i*_ = 2*p*_*i*_. For each locus, estimate *μ*_*i*_ as the observed mean of ***m***_*i*_, then compute 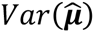 as the empirical variance across loci of 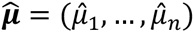, and because 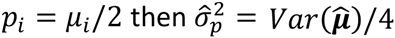 and 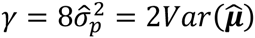.

##### 2-Considering pedigree structure

At locus *i*, gene content can be seen as a quantitative trait where the mean of ***m***_***i***_ in the base population is 2*p*_*i*_, where *p*_*i*_ is the allelic frequency at the base population, and the genetic variance is 2*p*_*i*_*q*_*i*_ [20]. Cockerham (1969) showed that the covariance of gene content of marker *i* across individuals *j* and *k* is a function of relationship *Cov*(*m*_*i,j*_, *m*_*i,k*_) = *A*_*jk*_2*p*_*i*_*q*_*i*_. A linear model can therefore be written as:

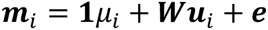

where ***W*** is an incidence matrix relating individuals in pedigree to genotypes, and with ***u***_***i***_ being the deviation of each individual from the mean *μ*_*i*_ for all individuals (Gengler et al., 2007; Forneris et al., 2015). Assuming multivariate normality:

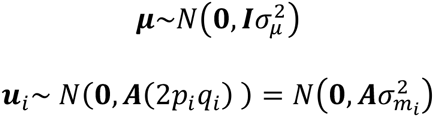

Equivalently, for the set of genotyped individuals (labelled as “2”), ***u***_2,*i*_~ *N*(**0, *A***_22_(2*p*_*i*_*q*_*i*_)) where ***A***_22_ = ***WAW*′** is an additive relationship matrix spanning only the genotyped individuals. From this formulation, there are two possible strategies to estimate 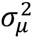.

<u>Generalized Least Squares (GLS)</u>. This ignores the prior distribution of ***μ*** and estimates each *μ*_*i*_ as a “fixed effect” using for each locus separate BLUP (or, equivalently, GLS) estimators of *μ*_*i*_. One option is to use the complete ***A***^−1^ and mixed model equations [20,21]. Equivalently, the corresponding GLS expression is

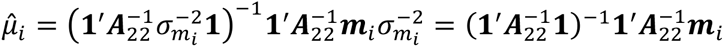

where (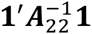) is the sum of elements of 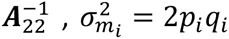 and 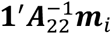 is simply a weighted sum of genotypes. Then, estimate 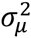 as 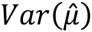 and because 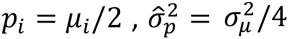, and it follows that 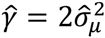.

<u>Maximum likelihood (ML).</u> Actually (and more exactly), *μ*_*i*_ can be considered as drawn from a normal distribution, 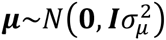. Thus 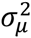 is a variance component that can be estimated by Maximum Likelihood. The equations for given values of 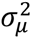 and 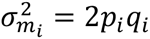 are 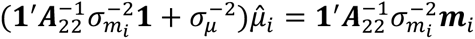. An Expectation-Maximization scheme [22] is as follows. Pick starting values for 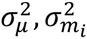. Iterate until convergence on:

1. For each marker *i*,

a. estimate 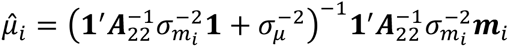
b. store 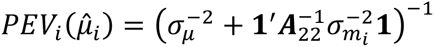
c. update 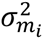 as 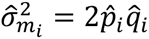 with 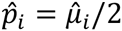
2. Update 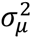 as 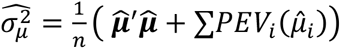, where the second part of the expression corresponds to the trace *Tr*(***IC***), ***I***, the identity matrix, is the relationship across ***μ*** and ***C*** is the prediction error covariance matrix of 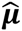. As only the diagonal elements of ***C*** are needed, the elements 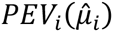 can be obtained separately from each single locus analysis.

On convergence, the estimate is 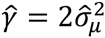. This gives the same estimate as the method based on a Wishart likelihood function in Christensen (2012) with *s* = *n*/2 (results not shown).

#### Estimation in multiple populations

If *t* base populations are considered, the variance component 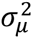 generalizes to ***Σ***_0_, a *t*×*t* matrix of variances and covariances across means 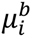 for marker *i* in population *b.* Across different populations, 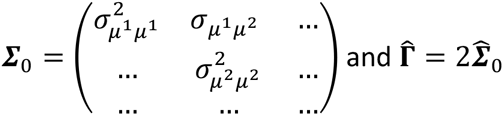.

##### 1-Assuming no pedigree structure

<u>NAIVE</u> If relationships across individuals are ignored:

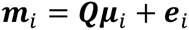

where ***Q*** is a matrix allocating individuals to populations and ***μ***_*i*_ is a vector with *t* elements including each population average. For each locus, ***μ***_*i*_ can be computed using least squares and the covariance matrix of ***μ***_*i*_ across loci gives an estimate of 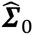.

##### 2-Considering pedigree structure

If there are no crosses, the estimation of allelic frequencies can be split in separate analysis by population 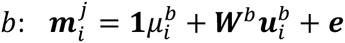 with 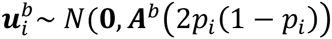, and ***A***^*b*^ is the matrix of relationships concerning population *b*. Then, 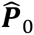 is estimated as the observed matrix of covariances across loci for estimated 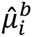. If there are crosses, there are two alternatives.

<u>GENERALIZED LEAST SQUARES (GLS).</u> The first alternative, suggested by Forneris et al. (2015) is to use a genetic groups model [11,23], as ***m***_*i*_ = ***Qμ***_*i*_ + ***Wu***_i_ + ***e*** where ***Q***_*k,b*_ contains the fraction of ancestry *b* in individual *k.* This ignores the fact that the variance of gene content, (2*p*_*i*_*q*_*i*_) is different for each breed and cross. The second, and more exact alternative is to use the representation where the breeding values are split into within and across breed components (Garcia-Cortes and Toro, 2006), as

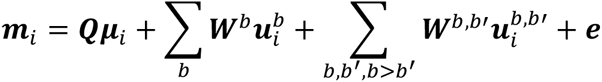

with partial relationship matrices for vectors ***u***^*b*^, ***u***^*b,b*^′.

<u>MAXIMUM LIKELIHOOD (ML).</u> Analogously to the single population case, an Expectation-Maximization updated estimate can be obtained using multiple trait formulations [22] where *PEC* is the prediction error variance-covariance, e.g. with two populations:

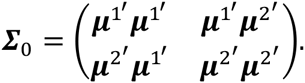

Our current implementation is as follows:

1. For each marker *i*,

a. estimate 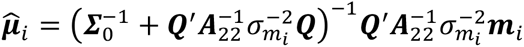
b. store 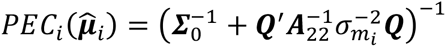
c. update 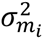 as 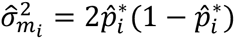 with 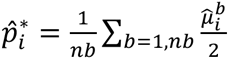
2. Update ***Σ***_0_ using crossproducts within and across populations as e.g. with two populations,

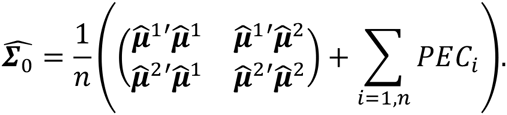

There is an approximation in (1c) because we assume that 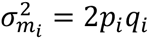 is equal across all base populations. This point will be addressed in future research.

## SIMULATION

To assess the quality of genomic predictions using one metafounder, we simulated data using QMSim [24]. The simulation closely followed Vitezica *et al.* (2011) to mimic a dairy cattle selection scheme scenario. A historical population undergoing mutation and drift was generated, followed by a recent population undergoing selection.

First, 100 generations of the historical population were generated with an effective population size of 100 during the first 95 generations, followed by a gradual expansion during the last 5 generations to an effective population size of 3000. In total 30 chromosomes of 100 cM and 40,000 segregating biallelic markers distributed at random along the chromosomes in the first generation of the historical population were simulated. The 40,000 markers were resampled from a larger set of 90,000 markers in order to obtain allelic frequencies from a beta(2,2) distribution, similar to dairy cattle marker data, so that true *γ* had a value around 0.40. Potentially, 1500 QTL affected the phenotype; QTL allele effects were sampled from a Gamma distribution with a shape parameter of 0.4. The mutation rate of the markers (recurrent mutation process) and QTL was assumed to be 2.5 × 10^−5^ per locus per generation (Solberg *et al.*, 2008). A female trait with a heritability of 0.30 was simulated.

Then, 10 overlapping generations of selection followed, where 200 males were mated with 2600 females producing 2600 offspring following a positive assortative mating design. Within the simulation, individuals were selected according to estimated breeding value (EBV) based on pedigree BLUP. In each generation 40% of the males and 20% of the females were replaced by younger and selected individuals. No restrictions were set to avoid or minimize inbreeding, so highly inbred individuals were found, as a result of extreme selection and matings among highly related individuals. There were 100 individuals (mainly found in the last generation) with an inbreeding coefficient higher than 0.20, with extreme cases (few individuals) with inbreeding coefficients higher than 0.40. True breeding values (TBV) and pedigree information were available for all 10 generations (28,800 individuals in pedigree), phenotypes were available for all females except the last generation (14,300 records). All males (840 sires of females with phenotypic records) were genotyped as well as 2600 individuals in generation 9 (with records) and 2600 in generation 10 (with no records). All in all, 20 independent replicates were made. A two-step analysis was carried out using the simulated data. First, we compared several methods to estimate *γ*. Then, we tested the quality of genomic predictions using four methods, one of them including one metafounder.

### Methods to estimate Gamma

Parameter *γ* was estimated using four different estimation methods. First, the NAIVE method which does not consider the pedigree structure. Then, the genealogical information was included in the estimation by three different methods: GLS, ML, and the Method of Moments (MM) presented in Legarra *et al.* (2015). For a single population, the last method involves the estimation of γ based on summary statistics of ***A***_22_ (regular pedigree-relationship matrix for genotyped individuals) and ***G*** (the genomic relationship matrix).

### Genomic prediction methods

Genetic merit of the selection candidates in generation 10 (genotyped and with no phenotype records) was estimated using four methods. The first one was the pedigree based BLUP (PBLUP) based on phenotype and pedigree information. The second method was Single-Step GBLUP (SSGBLUP) in which genomic information is also taken into account; this method used the correction of [25] and is the default method used in most practical applications [25,26]. However, the implementation that we used does not include inbreeding in the setup of ***A***^−1^ [27], although it does consider it in 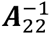 (see below for use of these matrices). The third method was Single-Step GBLUP including inbreeding in the setup of ***A***^−1^ and of 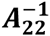 (SSGBLUP_F). Finally, the fourth method was SSGBLUP including the metafounder (SSGBLUP_M), using *γ* estimated by GLS as it turned out to be an accurate method to estimate gamma (see the Results section). The three methods used the following inverse relationship matrices: PBLUP: ***A***^−1^; SSGBLUP: 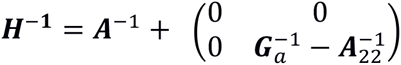 where ***G***_*a*_ is as in [25]; SSGBLUP_M: 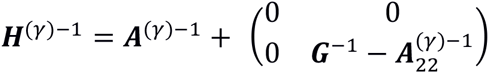 where ***G*** *=* (***M*** − ***J***)(***M*** − ***J***)’/*s* with *s* = *n*/2 (see the Methods section) and ***A***^(*γ*)^ is as in [1]. More details are given in Supplementary material. For computation we used blupf90 [28]. In the case of SSGBLUP_M we constructed all relationship matrices with own software, and then used the option user_file in blupf90.

### Quality of genomic prediction

Prediction quality was checked for all 2600 selection candidates. The accuracy of the methods was measured as the Pearson correlation between TBV and EBV. Bias was calculated as the difference between the average TBV and average EBV with respect to the base population. Thus, bias is related to estimated genetic progress in the selection candidates. The inflation (often called bias) of the prediction method was quantified by the coefficient of regression of TBV on EBV. These two statistics corresponds to the coefficients *b*_0_ and *b*_1_ in the Interbull validation method [29] which uses the regression *TSV* = *b*_0_ + *b*_1_*ESV* + *e*. The mean square error (MSE) was calculated as the mean of the squared difference between TBV and EBV. An ideal method should have maximum accuracy, minimum MSE, zero bias and a regression coefficient of 1. These are not only nice statistical properties but also have relevance in livestock selection [30–32]. Ranking changes of the selection candidates were also assessed by calculating the Spearman’s rank correlation coefficients between EBVs across methods.

In addition, the quality of variance component estimation was also assessed. For this purpose variance components were estimated using the four methods (PBLUP, SSGBLUP, SSGBLUP_F, SSGBLUP_M) using REML with remlf90 [28].

## RESULTS

### Estimation of gamma

Figure 1 shows boxplots of the differences between the estimates of *γ* calculated by four different methods (MM, Naive, ML and GLS) and the true values obtained by simulation, using each of the 20 replicates. The simulations were tailored to produce *γ* = 0.40. ML and GLS estimated *γ* very accurately. The MM clearly underestimated the value of *γ*, whereas the Naive method overestimated it. Based on these results we used the *γ* estimated by GLS when using SSGBLUP_M for prediction. The effect of employing different values of *γ* in the genomic prediction was assessed to quantify its impact in terms of the quality of predictions. Using estimates of *γ* based on the Method of Moments only slightly changed the results (not shown).

**Figure 1.**
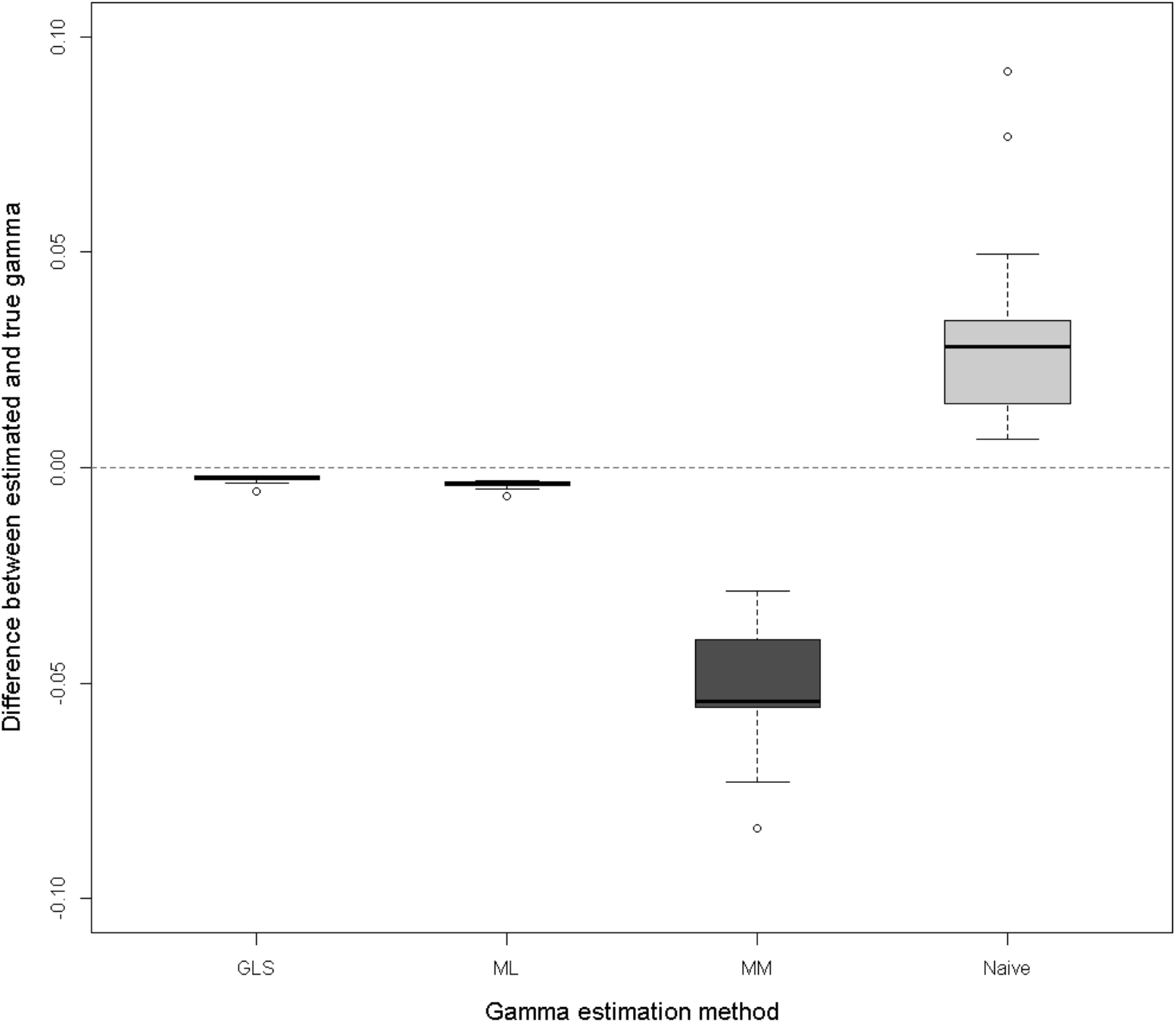
Differences between estimated and true Gamma, across 20 simulation replicates. Gamma was estimated by Generalized Least Squares (GLS), Maximum Likelihood (ML), Method of Moments (MM) and the Naive method.

### Quality of genomic prediction

Correlations between TBV and EBV for each of the prediction methods are shown in Table 1 and Figure 2a. Compared with PBLUP, SSGBLUP_F and SSGBLUP_M increased accuracy by approximately 23 absolute points, respectively. This shows an important improvement by including marker information in the prediction and the possibility of generating a small extra gain when also including the metafounder. SSGBLUP resulted in a small loss of accuracy as compared to SSGBLUP_F and SSGBLUP_M.

**Table 1.** **Accuracy (correlation between TBV and EBV), inflation (regression coefficient of TBV on EBV), bias (average (EBV-TBV)) and mean square error (MSE) for each of the prediction methods. Standard deviations in parenthesis.**

| Prediction method | Accuracy | Inflation | Bias | MSE |
| --- | --- | --- | --- | --- |
| PBLUP | 0.51 (0.05) | 0.98 (0.06) | -0.0003 (0.03) | 0.206 (0.01) |
| SSGBLUP | 0.72 (0.03) | 0.89 (0.19) | 0.2169 (0.04) | 0.159 (0.03) |
| SSGBLUP_F | 0.74 (0.02) | 0.99 (0.04) | 0.1167 (0.04) | 0.141 (0.01) |
| SSGBLUP_M | 0.74 (0.02) | 0.94 (0.04) | 0.0094 (0.03) | 0.125 (0.01) |

**Figure 2.**
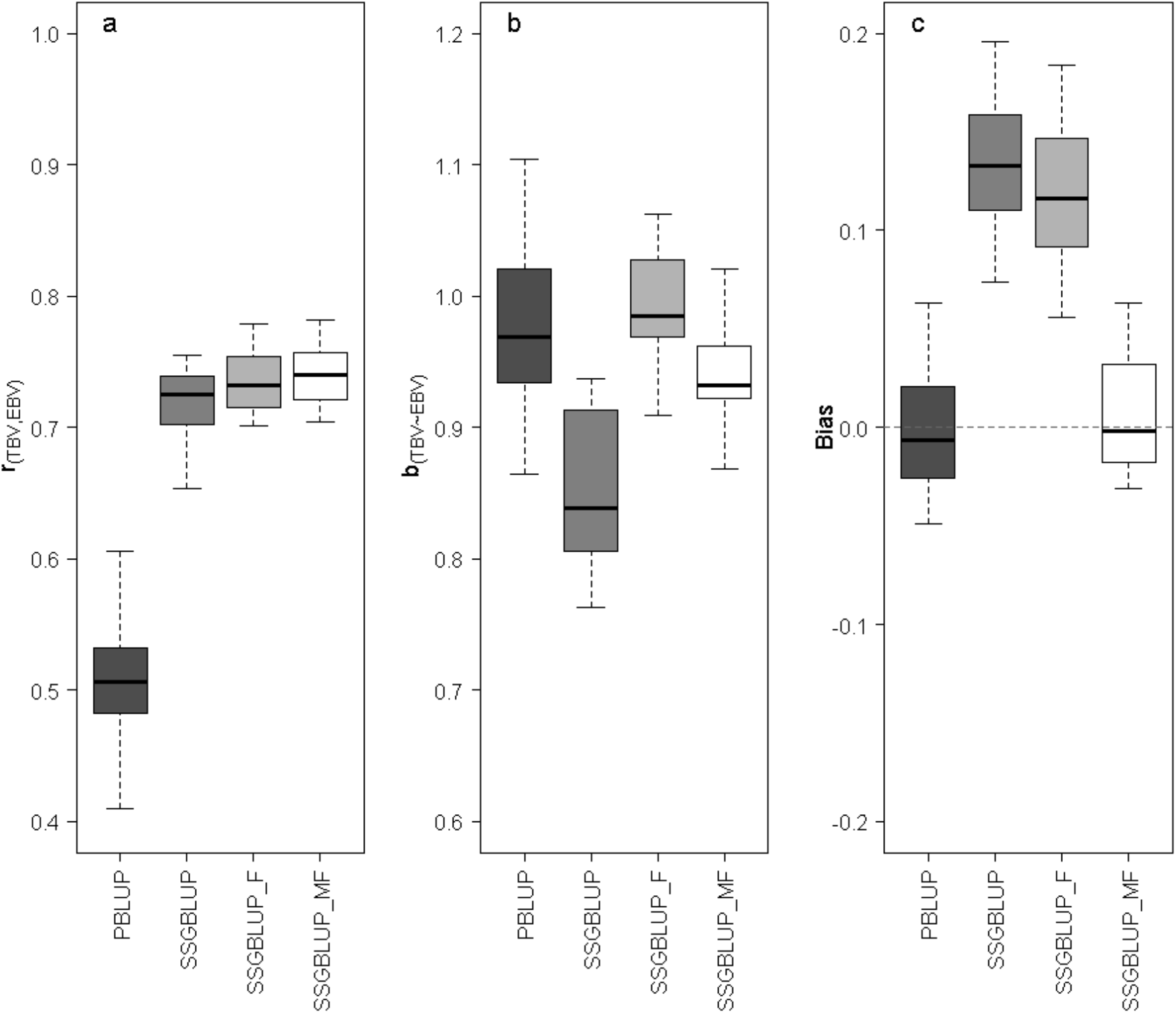
**a.** Correlation of TBV on EVB for each prediction method (accuracy). **b.** Regression slope of TBV on EBV (overdispersion). **c.** Bias (average (EBV-TBV)).

Bias values for each prediction method are shown in Table 1 and in Figure 2c. Both PBLUP and SSGBLUP_M were unbiased, whereas SSGBLUP and SSGBLUP_F were biased. Bias in SSGBLUP_F is equivalent to roughly 0.5 generations of genetic improvement or to 0.4 standard genetic deviations.

Table 1 and Figure 2b display the regression coefficient of TBV on EBV. This value measures the inflation degree of each prediction method and should be close to 1. PBLUP and SSGBLUP_F produced the values closest to one. Including genomic data in the prediction using SSGBLUP resulted in regression coefficients lower than one, but including the metafounder in SSGBLUP_M gives values closer to one. SSGBLUP_M and SSGBLUP_F displayed a lower standard deviation compared to the other two methods. Again, SSGBLUP showed the highest variability. SSGBLUP_M displayed the lowest MSE (closer to zero), followed by SSGBLUP_F (Table 1).

### Variance components estimation

Figure 3 shows the estimates of heritability obtained in three of the four methods assed (PBLUP, SSGBLUP_F and SSGBLUP_M). The estimates obtained using SSGBLUP are not displayed in Figure 3 because in 6 out of the 20 simulation replicates EM-REML did not converge. Convergence was achieved in those cases by weighting the submatrix 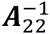 in ***H***^−1^ by *ω* = 0.7 instead of 1 [33] but poor quality estimates were obtained and they are not reported.

**Figure 3.**
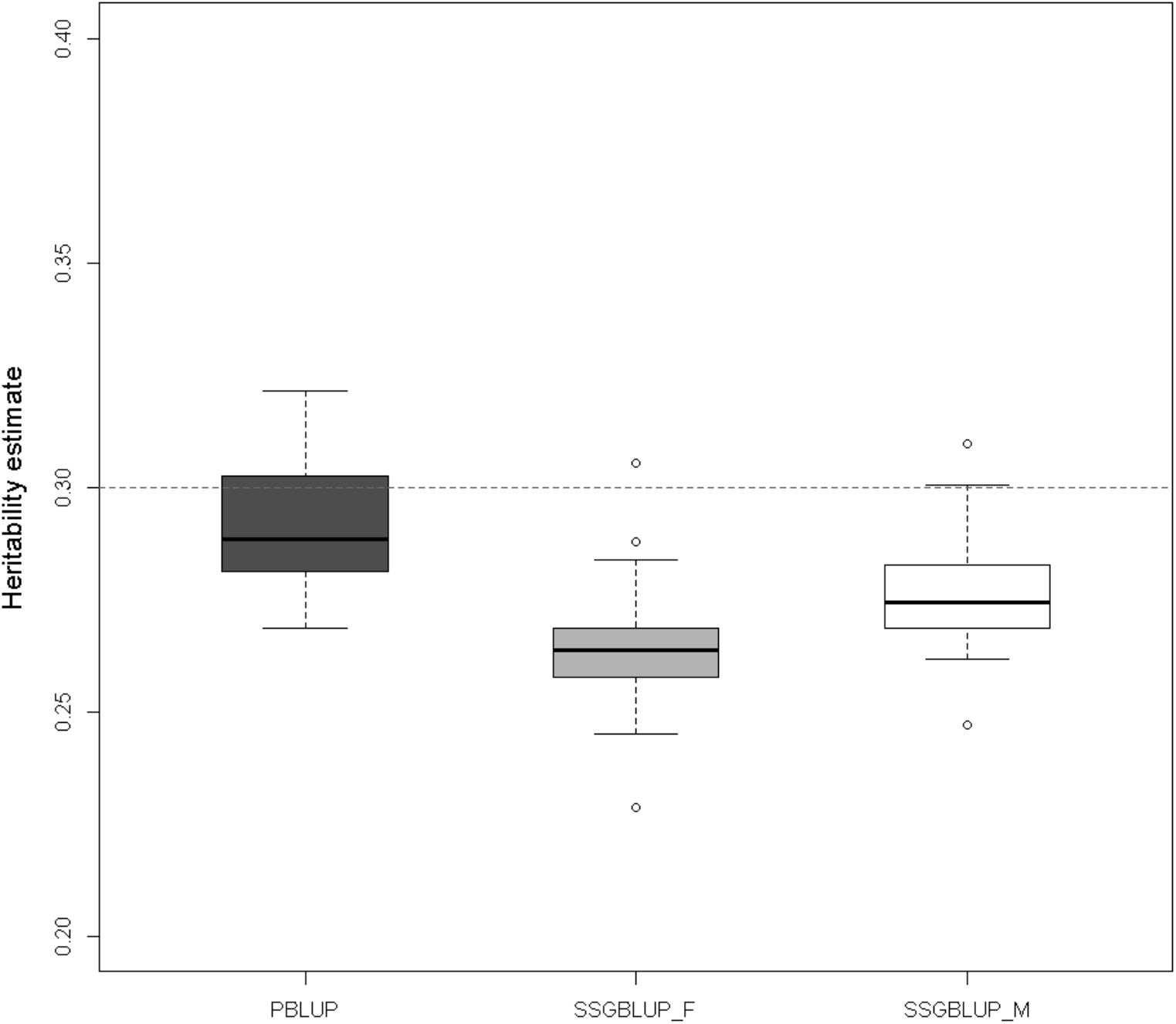
Estimated heritability for PBLUP, SSGBLUP_F and SSGBLUP_M considering the 20 replicates. The dotted line shows the simulated heritability of 0.30.

When comparing with the simulated true heritability value (0.30) the scenarios displayed in general lower estimates. The lowest estimates were obtained using SSGBLUP_F. Including the metafounder improved estimates compared to SSGBLUP_F and reduced variability when comparing to PBLUP.

### Ranking

The methods were also compared based on ranking correlations of EBVs with TBV and across methods. A rank correlation of 1 implies that the same candidates are selected. Results are in Table 2. Rank correlations with TBV are similar to accuracies in Table 1. Selection decisions are only slightly different using SSGBLUP, SSGBLUP_F or SSGBLUP_M. Note however, that this table does not address the comparison across generations (e.g. old vs. young animals), which is sensitive to biases reflected in Table 1 [32].

**Table 2.**
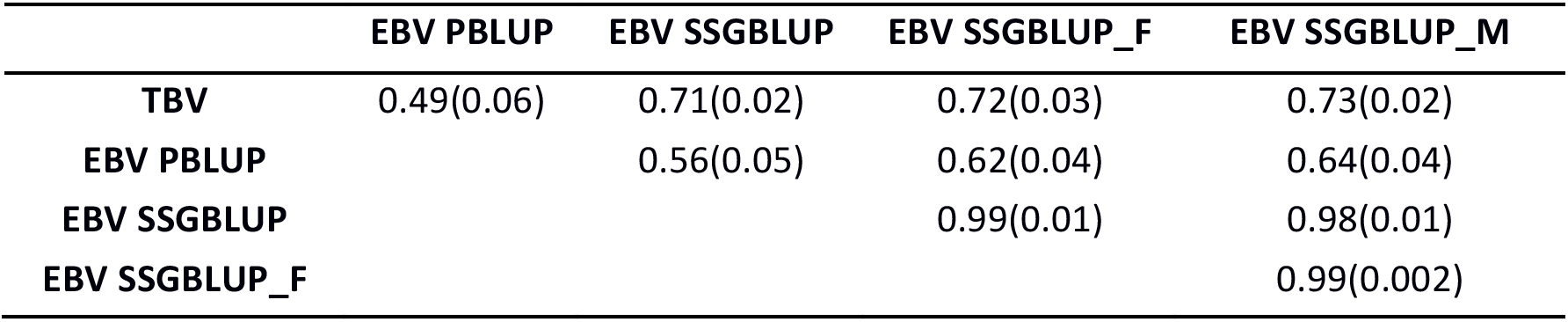
**Spearman correlation among TBV and the four EBV for each of the prediction methods. Standard deviations in parenthesis.**

|  | EBV PBLUP | EBV SSGBLUP | EBV SSGBLUP_F | EBV SSGBLUP_M |
| --- | --- | --- | --- | --- |
| **TBV** | 0.49(0.06) | 0.71(0.02) | 0.72(0.03) | 0.73(0.02) |
| **EBV PBLUP** |  | 0.56(0.05) | 0.62(0.04) | 0.64(0.04) |
| **EBV SSGBLUP** |  |  | 0.99(0.01) | 0.98(0.01) |
| **EBV SSGBLUP_F** |  |  |  | 0.99(0.002) |

## DISCUSSION

In this work, we have addressed the complex issue of conciliation of marker and pedigree information. Powell et al. [34] argued that both IBS (at the markers) and IBD are measures of identity at causal genes and they are compatible notions. However, the incompatibility issue appears when mixing both kind of relationships [5,25,35,36]. Legarra [7] established how to solve the issue of comparing genetic variance across IBD, IBS or other measures of relationships. In this work, we have used, similar (but not identical) to Powell et al. [34], a fixed reference (***G*** constructed as a crossproduct of {−1,0,1} genotypic codes) and tailored ***A*** (IBD, pedigree) to fit ***G*** (IBS, markers). Using a fixed reference has the advantage, compared to previous approaches, that genomic relationships are immutable (adding more genotypes to the database does not change the existing relationships) and they are unconditional on pedigree depth, that by construction is always limited and, in animal breeding, often heterogeneous. Our approach is in fact very similar to considering, as measures of identity, plain IBS. We use a matrix ***G*** = 2(***M*** − ***J***)(***M*** − ***J***)’/n, whereas a matrix of IBS, or molecular, relationships is ***G***_*IBS*_ *=* ***G***/2 + **11**′ (see proof at the supplementary material). In a GBLUP context when all animals are genotyped, using a model with IBS coefficients yields identical results as the term 1/2 gets absorbed into the variance component and the constant **11**′ gets absorbed into the fixed part of the linear mixed model [7,37]. However, the matrix that must be used in SSGBLUP_M is ***G*** and not ***G***_*IBS*_, because ***G***_*IBS*_ is not compatible with pedigree relationships.

## Easy estimation of ancestral relationships

The derivations in the THEORY section show that estimation of ancestral relationships in *γ* (one base population) and ***Γ*** (several base populations) may be framed within the linear model approach that is classical in quantitative genetics [13], and recently used for gene content [12,20,21]. These methods are easy to understand and to compute. Also, ***Γ*** can be understood, just like heritability, as an unobserved base population parameter that does not change with additional data (although its estimate may change). Therefore, an accurate estimate of ***Γ*** can be used repeatedly without the need of re-estimation, as is customary in livestock genetic evaluations. This contrasts with “centering” of marker covariates, which changes with every new genotype.

In the current research, the simplest methods (Naive and Method of Moments) yielded biased (upwards and downwards respectively) estimates of *γ*; for the first method because it ignores that allele frequencies drift to the extremes as generations go, and for the second because it implicitly assumes that individuals genotyped are a random sample from a particular generation when in fact they are not.

In addition, the equivalence of ancestral relationships with second moments of allele frequencies shows a strong relation with populations genetics theory, which will be detailed in the next paragraph.

## Relationship between metafounders γ and F_st_ fixation index

The fixation index *F*_*st*_ [38] is a measure of diversity of a set of populations with respect to a reference population, usually the pool of all populations. In this view, each population is a random sample from all possible populations that could be sampled according to the evolutionary process described by *F*_*st*_. Conceptually, *F*_*st*_ is a parameter to be estimated [13,39], and it is not a statistic computed from the data. A usual definition of *F*_*st*_ for a particular biallelic locus is

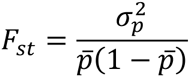

where 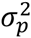 is the variance of allelic frequencies across populations and 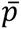 is the allelic frequency of the conceptual combined population. If we consider that the variance of allelic frequencies applies *across* loci and not *across* populations, it follows naturally that 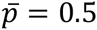. In this case,

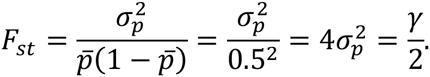

Our interpretation is as follows. Jacquard (1974) called 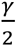 the “inbreeding coefficient of a population”. Cockerham (1969) modelled 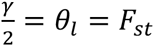 as an intraclass correlation, “the coancestry of the line with itself”, in other words, the probability that two gametes taken at random from the line are identical. Thus, it makes perfect sense to consider that the additive relationship (which is twice the coancestry value) of a group with itself is 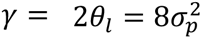. This is the interpretation of the 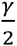 coefficient in Legarra et al. [1]. Note that the assumption 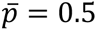 is automatically fulfilled if reference alleles are labelled randomly across loci (i.e., they are neither the most frequent nor the least observed).

Alternatively, Legarra et al. (2015) showed that for a population with self-relationship of *γ*, the average heterozygosity was 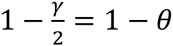, i.e. the variance is reduced by an amount of *θ* from the conceptual population with heterozygosity 1. Thus 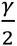 can be interpreted as *F*_*st*_ if the latter is taken as a measure of homozygosity.

## Consequences of using metafounders in genomic evaluation

Genomic estimates of breeding values are invariant to allele coding [37] when all individuals are genotyped. However, this is not the case when pedigree and marker information are combined as in SSGBLUP. In this work we have shown that, even in presence of complete pedigree and a single base population, use of metafounders in SSGBLUP_M leads to slightly more inflated, less biased EBVs, lower MSE and nearly unbiased estimates of heritability compared to SSGBLUP_F. Bias, defined as E(EBV-TBV)), is typically overlooked in genomic predictions, but in an example of biased evaluation “sires of later generations appeared to be under-evaluated relative to older sires” [40]. Overdispersion, also called bias in recent literature (e.g. Mantyssari et al., 2010), may have dramatical impact as well [30–32]. The trade-off between bias and variance needs further studies. For instance, [5] found that SSGBLUP_F was unbiased but had some overdispersion; this is likely dependent on the data structure, including the genotyping.

In addition, use of metafounders allows a clear definition of genomic relationships. With this definition, relationships are not dependent on pedigree depth or completeness, and are not dependent on allelic frequencies subject to change with arrival of new data. Additionally, a high dimensional parameter (-base- allele frequencies) is substituted by a low-dimensional one (matrix ***Γ***).

The poor performance of SSGBLUP as compared to SSGBLUP_F (the former ignoring inbreeding in the set up of **A**^−1^) is likely due to the presence of highly inbred individuals. This relates to the interpretation of an *ω* parameter used in early studies of SSGBLUP. An application of SSGBLUP for type traits in Holstein [33] experienced convergence problems. The authors found that by multiplying 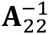 by a *ω* = 0.7 eliminated convergence problems and increased accuracy. However, the nature of that parameter was not known, e.g. Misztal et al. [41]. In those studies, the inverse of the numerator relationship matrix ***A***^−1^ was constructed using Henderson’s rules while ignoring inbreeding [27], while the submatrix 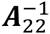 included inbreeding. Subsequently, the elements in the latter were too large. In addition, genotyped animals were on average unrelated in ***G*** but not in ***A***_22_, which is corrected by scaling ***G*** as in Vitezica et al. (2011). But then, in 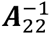 the elements were too large for younger animals relative to ***G***. Both problems are partially circumvented but putting a weight *ω* < 1 on 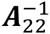 When ***A***^−1^ was constructed considering inbreeding, the optimal *ω* coefficient in an analysis of Holstein dairy cattle increased from 0.7 to 0.9 (Masuda, personal communication, 2016). However, the metafounder approach provides a clean solution to this problem. Also, following these experiences, ***A***^−1^ should always be constructed considering inbreeding to avoid pathological problems.

## CONCLUSION

Metafounders are similar to F_st_ fixation indices and proportional to covariances of allelic frequencies in base populations. Use of metafounders is simplified by new methods (GLS and maximum likelihood) to estimate the covariance of base allele frequencies. We verified by simulation of a selected population that, in a single population, both GLS and ML are unbiased and computationally efficient. In the same simulation, use of metafounders in Single Step GBLUP leads to more accurate and less biased evaluations, and also to more accurate estimates of genetic parameters.

We propose a genomic relationship matrix that refers to a population with ideal frequencies 0.5. This matrix is similar to an IBS relationship matrix (up to scale factors), does not change with new data and is compatible with pedigree data if metafounders are used.

In this simulated data, pedigrees are perfectly known. Future work with real data sets in more complex settings - purebreds and their crosses [42,43], and selected populations with unknown parent groups [11] will investigate the feasibility and accuracy in practice of using metafounders on Single Step GBLUP.

## APPENDIX

This Appendix contains several algebraic developments not detailed in the main text.

### Analytical derivation of *γ* and *s*

For a particular population, the genetic variance-covariance structure is a function of two parameters *η*_1_ and 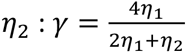 and *s* = *n*(2*η*_1_ + *η*_2_) (*n* being the number of markers) which depend on the allelic frequencies (Christensen 2012), Appendix A. With *p*_*j*_ being the allelic frequencies across the *j* = 1. *n* loci, these parameters do not depend on *j* and are equal to

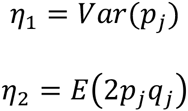

with *q* = 1 − *p*.

Now use is made of the following developments.

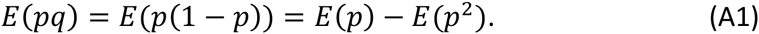

Since we have that *Var*(*p*) = *E*(*p*^2^) − *E*(*p*)^2^ we obtain *E*(*p*^2^) = *Var*(*p*) + *E*(*p*)^2^. We also have *E*(*q*) = 1 − *E*(*p*). Substituting *E*(*p*)^2^ in (A1) gives

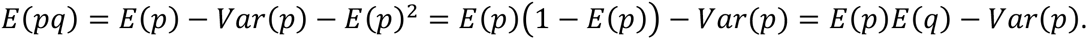

If markers are biallelic and labelled at random *E*(*p*) = *E*(*q*) = 0.5. So the equation above gives *E*(*pq*) = 0.25 − *Var*(*p*). From this we obtain

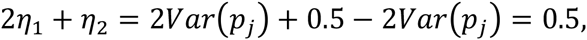

and therefore

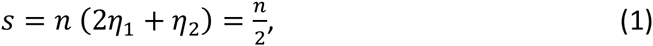

or, in other words, *s* is half the number of markers. Further,

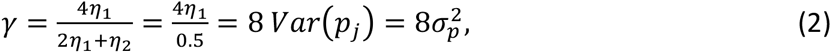

so that *γ* for a single population is eight times the variance of allelic frequencies at the base population.

### Equivalences of genomic relationship matrices

The matrix **G** described in Christensen (2012) and in this paper can be written as 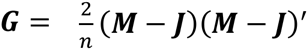, where **M** contains genotypes coded as {0,1,2} and ***J*** is a matrix of 1’s. The purpose of this paragraph is to show the linear relationship of this matrix with a matrix describing identity by state coefficients (IBS), in fact 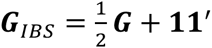. The terms in ***G***_*IBS*_ are usually described in terms of identities or countings (i.e. Ritland, 1996; Toro et al., 2011; Nejati-Javaremi et al., 1997):

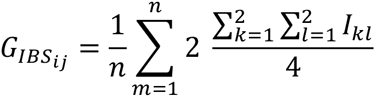

where *I*_*kl*_ measures the identity (with value 1 or 0) of allele *k* in individual *i* with allele *l* in individual *j*, and single-locus identity measures are averaged across *n* loci. There is an algebraic expression for this “counting”. Toro et al. (2011) expression (1), show that for biallelic markers, for a locus k (omitted in the notation for clarity):

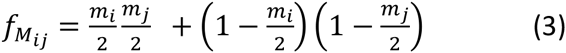

for coancestry (half relationship) 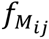 of individuals i and j, where *m*/2 is the “gene frequency” of the individual (half *m* the gene content, i.e. {0,1/2,1} for the three genotypes).

In order to prove 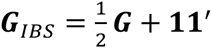, first we translate the Toro et al. (2011) equation to the more familiar scale of relationships 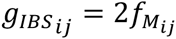 and gene contents *m*. Thus

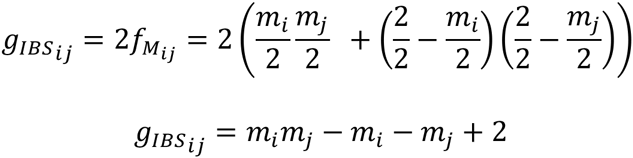

This expression can be easily verified in a table with the nine possible genotypes:

|  | AA | Aa | aa |
| --- | --- | --- | --- |
| AA | 2 | 1 | 0 |
| Aa | 1 | 1 | 1 |
| aa | 0 | 1 | 2 |

Also,

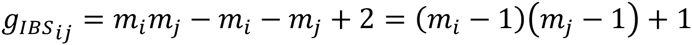

which extends to all individuals and averaged across loci can be written as

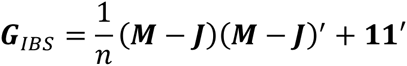

Thus, matrix 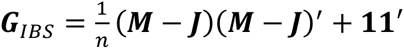 and because 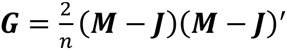 it follows that 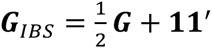. The equivalence can also be verified by noting that, for all nine genotypes, the cross-product (*m*_*i*_ − 1) in the following table is identical to 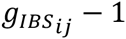 in the previous table.

|  | AA | Aa | aa |
| --- | --- | --- | --- |
| AA | 1 | 0 | -1 |
| Aa | 0 | 0 | 0 |
| aa | -1 | 0 | 1 |

### Computation of the different H matrices

For SSGBLUP and SSGBLUP_F, matrix ***H***^−1^ is constructed as follows:

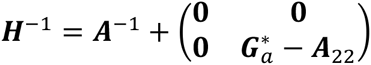

with 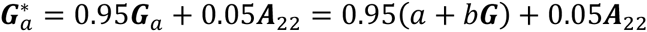, and 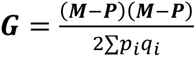 as in VanRaden (2008), **M** contains genotypes coded as {0,1,2} and **P** contains twice allelic frequencies ***p***_***i***_. These are computed from the observed genotypes so that 2*p*_*i*_ is equal to the the mean of the *i*-th column of **M**. Constants *a* and *b* are such that the full-matrix and diagonal averages of ***G***_*a*_ and ***A***_22_ are the same (Christensen et al., 2012) in order to make the two matrices compatible. The use of the weights 0.95 and 0.05 is in order to make ***G***_*a*_ invertible. Matrix ***A***^−1^ should be constructed using contributions with values described in the Table below (i.e. Meuwissen and Luo, 1992):

|  |  |
| --- | --- |
| No parent known | 1 |
| One parent known | $\left(0.75 - \frac{F_{known}}{4}\right)^{-1}$ |
| Two parents known | $\left(0.5 - \frac{F_{sire}}{4} - \frac{F_{dam}}{4}\right)^{-1}$ |

Or, in a more compact way 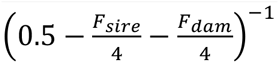 with *F*_*unknown*_ = −1.

SSGBLUP uses the defaults in blupf90 suite of programs (random_type*add_animal*). SSGBLUP uses the simple method to create

***A***^−1^, method which pretends that in all cases inbreeding in expressions above is *F* = 0.

SSGBLUP_F uses ***H***^−1^ as above but constructs ***A***^−1^ correctly (blupf90 random_type *add_an_upginb*), using the rules above.

SSGBLUP_M uses the blupf90 random_type *user_file* to consider the following relationship matrix:

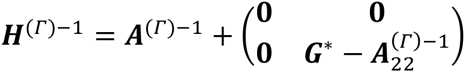

with 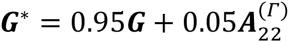 (basically to make ***G*** invertible), 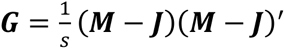 and *s* = *n*/2, **M** contains genotypes coded as {0,1,2}, *n* is the number of markers, ***A***^(*Г*)−1^ and 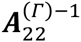 are constructed with own programs as in Legarra et al. (2015) using the estimated value of ***Γ***. Inbreeding is fully considered in both matrices.

## DECLARATIONS

Availability of data and materials: Software and files are avaible at https://github.com/alegarra/metafounders.

## Authors contribution

AL and OFC derived the theory with help from ZGV and CAGB. All authors agreed on scenarios to be tested. CAGB programmed and run all the simulations, with substantial input from IP and IM. The initial version of the manuscript was written by CAGB and AL and then completed by all authors.

## Acknowledgments

We thank S Boitard and B Servin for discussions concerning Fst and all members of AdMixSel project.

## REFERENCES

1 Legarra A, Christensen OF, Vitezica ZG, Aguilar I, Misztal I. Ancestral Relationships Using Metafounders: Finite Ancestral Populations and Across Population Relationships. Genetics. 2015;200:455–68.

2 Legarra A, Aguilar I, Misztal I. A relationship matrix including full pedigree and genomic information. J Dairy Sci. 2009;92:4656–63.

3 Christensen OF, Lund MS. Genomic prediction when some animals are not genotyped. Genet Sel Evol. 2010;42:2.

4 Fernando RL, Dekkers JC, Garrick DJ. A class of Bayesian methods to combine large numbers of genotyped and non-genotyped animals for whole-genome analyses. Genet Sel Evol. 2014;46:50.

5 Vitezica Z, Aguilar I, Misztal I, Legarra A. Bias in genomic predictions for populations under selection. Genet. Res. 2011;93:357–66.

6 Christensen OF, Legarra A, Lund MS, Su G. Genetic evaluation for three-way crossbreeding. Genet. Sel. Evol. 2015;47:98.

7 Legarra A. Comparing estimates of genetic variance across different relationship models. Theor. Popul. Biol. 2016;107:26–30.

8 VanRaden PM. Efficient Methods to Compute Genomic Predictions. J Dairy Sci. 2008;91:4414–23.

9 Ritland K. Estimators for pairwise relatedness and individual inbreeding coefficients. Genet. Res. 1996;67:175–85.

10 Christensen OF. Compatibility of pedigree-based and marker-based relationship matrices for single-step genetic evaluation. Genet. Sel. Evol. 2012;44:37.

11 Quaas RL. Additive genetic model with groups and relationships. J. Dairy Sci. 1988;71:1338–45.

12 Makgahlela M, Strandén I, Nielsen U, Sillanpää M, Mäntysaari E. Using the unified relationship matrix adjusted by breed-wise allele frequencies in genomic evaluation of a multibreed population. J. Dairy Sci. 2014;97:1117–27.

13 Cockerham CC. Variance of gene frequencies. Evolution. 1969;23:72–84.

14 Wright S. Evolution in Mendelian populations. Genetics. 1931;16:97–159.

15 Crow J, Kimura M. An introduction to population genetics theory. Harper and Row, New York; 1970.

16 Toro MÁ, García-Cortés LA, Legarra A. A note on the rationale for estimating genealogical coancestry from molecular markers. Genet. Sel. Evol. GSE. 2011;43:27.

17 Robertson A. Gene Frequency Distributions as a Test of Selective Neutrality. Genetics. 1975;81:775–85.

18 Fariello MI, Boitard S, Naya H, SanCristobal M, Servin B. Detecting signatures of selection through haplotype differentiation among hierarchically structured populations. Genetics. 2013;193:929–941.

19 Laval G, SanCristobal M, Chevalet C. Measuring genetic distances between breeds: use of some distances in various short term evolution models. Genet. Sel. Evol. 2002;34:481–508.

20 Forneris NS, Legarra A, Vitezica ZG, Tsuruta S, Aguilar I, Misztal I, et al. Quality control of genotypes using heritability estimates of gene content at the marker. Genetics. 2015;199:675–81.

21 Gengler N, Mayeres P, Szydlowski M. A simple method to approximate gene content in large pedigree populations: application to the myostatin gene in dual-purpose Belgian Blue cattle. animal. 2007;1:21–8.

22 Mäntysaari E, Vleck L. Restricted maximum likelihood estimates of variance components from multitrait sire models with large number of fixed effects. J. Anim. Breed. Genet. 1989;106:409–22.

23 Thompson R. Sire evaluation. Biometrics. 1979;35:339–53.

24 Sargolzaei M, Schenkel FS. QMSim: a large-scale genome simulator for livestock. Bioinformatics. 2009;25:680–1.

25 Christensen O, Madsen P, Nielsen B, Ostersen T, Su G. Single-step methods for genomic evaluation in pigs. Animal. 2012;6:1565–71.

26 Masuda Y, Misztal I, Tsuruta S, Legarra A, Aguilar I, Lourenco DAL, et al. Implementation of genomic recursions in single-step genomic best linear unbiased predictor for US Holsteins with a large number of genotyped animals. J. Dairy Sci. 2016;99:1968–1974.

27 Mehrabani-Yeganeh H, Gibson JP, Schaeffer LR. Including coefficients of inbreeding in BLUP evaluation and its effect on response to selection. J. Anim. Breed. Genet. 2000;117:145–51.

28 Misztal I, Tsuruta S, Strabel T, Auvray B, Druet T, Lee DH. BLUPF90 and related programs (BGF90). 7th World Congr. Genet. Appl. Livest. Prod. Montpellier, France; 2002. p. CD-ROM Communication N° 28–07.

29 Mantysaari E, Liu Z, VanRaden P. Interbull validation test for genomic evaluations. Interbull Bull. 2010;41.

30 Sargolzaei M, Chesnais J, Schenkel FS. Assessing the bias in top GPA bulls [Internet]. 2012 [cited 2016 Jul 21]. Available from: cgil.uoguelph.ca/dcbgc/Agenda1209/DCBGC1209_Bias_Mehdi.pdf

31 Spelman RJ, Arias J, Keehan MD, Obolonkin V, Winkelman AM, Johnson DL, et al. Application of genomic selection in the New Zealand dairy cattle industry. Proc. 9th World Congr. Genet. Appl. Livest. Prod. 1-6 August 2010 Leipz. [Internet]. 2010 [cited 2016 Jul 26]. Available from: http://www.icar.org/Cork_2012/Manuscripts/Published/Spelman.pdf

32 Winkelman AM, Johnson DL, Harris BL. Application of genomic evaluation to dairy cattle in New Zealand. J. Dairy Sci. 2015;98:659–75.

33 Tsuruta S, Misztal I, Aguilar I, Lawlor T. Multiple-trait genomic evaluation of linear type traits using genomic and phenotypic data in US Holsteins. J. Dairy Sci. 2011;94:4198–204.

34 Powell JE, Visscher PM, Goddard ME. Reconciling the analysis of IBD and IBS in complex trait studies. Nat Rev Genet. 2010;11:800–5.

35 Harris BL, Johnson DL. Genomic predictions for New Zealand dairy bulls and integration with national genetic evaluation. J Dairy Sci. 2010;93:1243–52.

36 Meuwissen T, Luan T, Woolliams J. The unified approach to the use of genomic and pedigree information in genomic evaluations revisited. J. Anim. Breed. Genet. 2011;128:429–39.

37 Strandén I, Christensen OF. Allele coding in genomic evaluation. Genet Sel Evol. 2011;43:25.

38 Wright S. Isolation by Distance. Genetics. 1943;28:114–38.

39 Holsinger KE, Weir BS. Genetics in geographically structured populations: defining, estimating and interpreting FST. Nat. Rev. Genet. 2009;10:639–650.

40 Henderson CR. Sire evaluations and genetic trends. J Anim Sci. 1973;Symposium.

41 Misztal I, Vitezica Z-G, Legarra A, Aguilar I, Swan A. Unknown-parent groups in single-step genomic evaluation. J. Anim. Breed. Genet. 2013;130:252–8.

42 Christensen OF, Madsen P, Nielsen B, Su G. Genomic evaluation of both purebred and crossbred performances. Genet. Sel. Evol. 2014;46:1–9.

43 Lourenco DAL, Tsuruta S, Fragomeni BO, Chen CY, Herring WO, Misztal I. Crossbreed evaluations in single-step genomic best linear unbiased predictor using adjusted realized relationship matrices. J. Anim. Sci. 2016;94:909.

